# Livestock heat stress risk in response to the extreme heat event (heatwave) of July 2022 in the UK

**DOI:** 10.1101/2023.05.18.541284

**Authors:** A. S. Cooke, M. J. Rivero

## Abstract

On the 18^th^ and 19^th^ of July 2022, the UK experienced a record-breaking extreme heat event. For the first time, temperatures exceeding 40°C were recorded. Whilst this may seem exceptional or unprecedented, the progression of climate change is expected to increase both the likelihood and severity of such events. Livestock are vulnerable to heat stress, which manifests as losses to health and welfare, productivity, and sustainability. Here, we characterize the heatwave of July 2022 in the context of livestock heat-stress risk, with a focus on cattle. Meteorological data was obtained from 85 weather stations and the Comprehensive Climate Index (CCI) was calculated, hourly, for each station. The CCI was mapped across the UK for 18/07/22 and 19/07/22 and compared against heat stress risk thresholds. Across both days, >25% of sites experienced “severe” heat stress risk. On 19/07/22 there was an “extreme” risk across >5% of sites. The site that experienced the highest risk was near Rugby, in the West Midlands. Across all sites, night-time temperatures fell below risk thresholds and may have mitigated some of the heat stress risk. Whilst there was some evidence of productivity losses, this was not conclusive. The impacts of this event on livestock were not just direct, but indirect through negative impacts on water and forage availability. The heatwave of July 2022 must serve as a warning for the UK livestock industry and these results may act as a case study of what the sector may be increasingly likely to experience in the future.

## 1 Introduction

In the last decades, livestock species have been severely affected by heat stress because of increasing temperatures, which has threatened animal welfare and decreased production (Carvajal et al., 2021); dairy cows produce less milk with lower milk quality characteristics, whilst in beef cattle, heat stress impairs reproductive performance of nursing cows, decreases growth rate, and worsens meat quality in growing/finishing animals (Summer et al., 2019). Actually, ca. 7% of the global cattle population is currently exposed to dangerous heat conditions, and this percentage is projected to increase to ∼48% before 2100 under a scenario of growing emissions, being poor and livestock-dependent tropical countries the most affected (Carvajal et al., 2021). In the Northern Hemisphere, the most severe heat stress is expected during the months of July and August, since in many instances the temperature does not drop enough to allow the animals to completely dissipate heat gained during the preceding day.

In July 2022 the UK and much of Europe experienced an extreme heat event (heat wave), with air temperature exceeding 40°C in some areas, setting new records as well as a new national record for the hottest temperature recorded in the UK of 40.3 °C at Coningsby in Lincolnshire. For the first time ever, for the days of July 18^th^ and 19^th^, the Met Office issued a ‘Red Weather Warning’ for heat, meaning “*dangerous weather*” and “*risk to* [human] *life*” (Met Office, 2022). Whilst such temperatures may be commonplace across much of the world, they are not in the UK. Consequently, UK livestock are not physiologically adapted or acclimatised to such extremes and not necessarily are livestock systems. Indeed, adaptation for cold weather has arguably been preferable. Furthermore, such events are predicted to be more frequent and more severe due to the impacts of climate change (IPCC, 2022).

Heat waves (consecutive days of severe or extreme heat) can cause heat stress events thus reducing animal performance and leading to welfare, economic, and environmental losses in livestock systems (Dunn et al., 2014; Garner et al., 2017; Lees et al., 2019). The effect of these extreme conditions can be easily verified on dairy cattle since the monitoring of daily records of milk production can quickly identify any drop in yield with and the associated immediate effect on the income generated. However, for beef cattle or lamb, the detrimental effect of heat stress can take longer to be identified, e.g., between two consecutive weighing events. Actually, one of the most popular heat stress indices, i.e., the Temperature-Humidity Index or THI, had its earliest example of application as the basis for livestock response functions for milk production decline of dairy cows in 1964 (Berry et al., 1964; Hahn et al., 2009).

The THI has for several years served as a de facto standard for classifying thermal environments in many livestock production and transport situations, and a basis for strategic and tactical management practices during seasons other than winter (Hahn et al, 2009). Modifications to the THI have been proposed to overcome limitations related to lack of inclusion of airflow and radiation heat loads (Mader et al., 2006). To overcome these limitations, Mader et al., (2010) developed the Comprehensive Climate Index (CCI) that incorporates major environmental components that are experienced over a range of hot and cold conditions and established environmental stress thresholds reflecting stress levels based on environmental conditions, management levels, and physiological status. CCI also works on a Celsius basis as opposed to THI which works off of Fahrenheit. This study aimed to characterise the extent and spatial and temporal nature of the extreme heat event that occurred in the UK on 18 and 19 July 2022 in the context of livestock heat stress risk.

## 2 Methods

Meteorological data was taken from the Met Office Integrated Data Archive System (MIDAS) network, accessed via the Centre for Environmental Data Analysis (CEDA) (Met Office, 2012). Stations were selected if they met both of two criteria: (1) recorded data for all of the four weather variables of air temperature, relative humidity, windspeed, and solar radiation (2) were on mainland Great Britain (inc. Anglesey) or Northern Ireland. A total of 85 stations met these criteria (Figure 1). Individual stations were identified by their source ID (SRC_ID) as per the MIDAS database.

**Figure 1.**
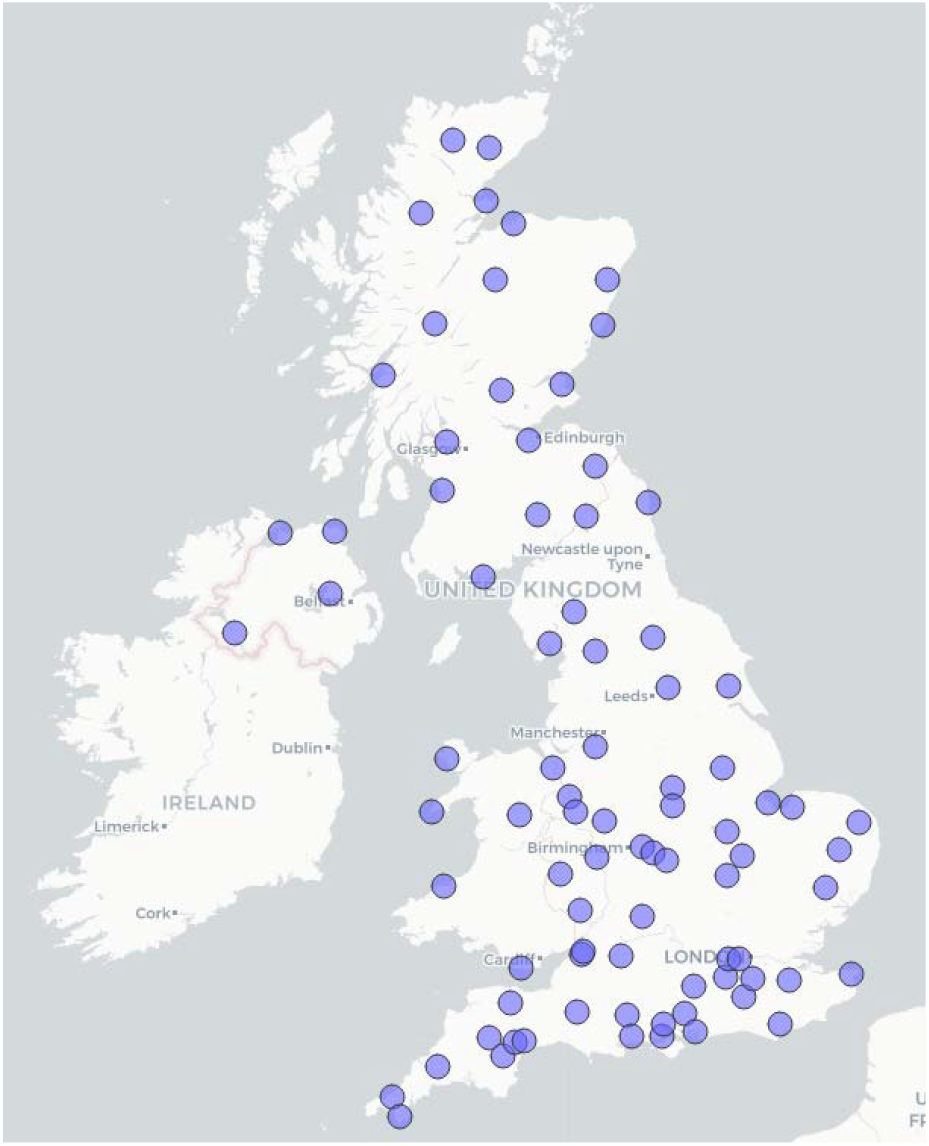
Location of MIDAS weather stations used in this study.

Hourly readings of the four weather variables to calculate an hourly CCI score (calculations as per Mader et al. (2010)) per station were used (i.e., air temperature, relative humidity, wind speed, solar radiation). MIDAS reports solar radiation in kilojoules per square metre for the hour, these values were divided by 3.6 to give Watts per square metre, as necessary for the CCI calculation. Wind speed was also converted from knots to metres per second. In 2017 one station (Londonderry SRC_ID 56963) had thirteen records of negative solar radiation values, which were removed. If readings were not present for all of the four required variables, the record for that time point for that station was removed. CCI values could then be compared to heat risk thresholds taken from Mader et al. (2010) (Table 1). Rainfall data was also obtained to compare to previous years.

**Table 1.** Arbitrary comprehensive climate index thermal stress thresholds. With severe thresholds capable of causing death of animals and extreme thresholds having a high probability of causing death of high-risk animals. Adapted from Mader et al. (2010).

| Heat risk | Threshold (CCI) |
| --- | --- |
| Extreme danger | > 45 |
| Extreme | > 40 – 45 |
| Severe | > 35 – 40 |
| Moderate | > 30 – 35 |
| Mild | > 25 – 30 |
| No stress | < 25 |

Heat maps were created for each of the two days from the hour with the highest mean national CCI values. Spatial interpolation for the maps was performed using the Inverse Distance Weighting (IDW) technique. For the period 16/07/22 to 21/07/22 (heat event ± two days) national hourly CCI figures were graphed showing the 50^th^ percentile (mean), 75^th^ percentile (3^rd^ quartile) and 95^th^ percentile. Additionally, the station with the highest mean CCI across the two days was plotted. Air temperature and CCI were directly compared across the extreme heat event to investigate the extent of differences between the two measures. For each individual component of CCI a comparison was made (using midday readings) between the extreme heat event of 2022 (18/07/22 to 19/07/22) and, for each previously July of 2017-2021, the two consecutive days with the highest CCI averaged across all the met stations.

For the purposes of contextualising the wider implications of the extreme heat event on the livestock industry, national slaughter data and milk data were obtained from Department for Environment, Food and Rural Affairs (DEFRA, 2022a, 2022b) and on-farm cattle deaths obtained via a request to the Rural Payments Agency under the Environmental Information Regulations (2004), equivalent data for sheep was unavailable as reporting of individual sheep deaths is not required in law. To illustrate the aspect of the ground cover prior, during and after the heatwave, satellite imagery and Normalised Difference Vegetation Index (NDVI) were taken from NASA (NASA, 2022).

### 2.1.1 Software

Heat maps were created using QGIS 3.26.1 (QGIS, 2022). Other figures were created in R Studio 1.2.1335 (running of R 4.2.0) using packages ‘ggplot2’ and ‘Cairo’ (R Core Team, 2021; R Studio Team, 2020; Urbanek and Horner, 2020; Wickham, 2016).

## 3 Results

Over the July periods of six years analysed (2017 to 2022), 99 of the 100 highest air temperatures were recorded occurred on 18/07/22 or 19/07/22, with the greatest being 40.0°C in Lincolnshire (SRC_ID 384) at 16:00 on 19/07/22 (this differs from widely publicised records due to different temporal resolutions). CCI values gave a similar, albeit less extreme, result, with 54 of the top 100 values being recorded on 18/07/22 or 19/07/22.

July 2022 also yielded particularly low rainfall, with a national mean of 48.4mm, the lowest since 1999. From 01/07/22 to 18/07/22 mean total rainfall was 19.0mm, thus the majority of rain occurred after the heat wave. Daily mean rainfall across the UK was 0.088 on both 18/07/22 and 19/07/22, with 90.7% of MIDAS weather stations recording no rain on the 18^th^ and 95.0% recording no rainfall on the 19^th^.

Both days with the Red Weather Warning showed high CCI scores across the country, particular for southern and eastern regions (Figure 2). Whilst CCI did reduce in some western areas on the second day, this was also when levels peaked elsewhere. This resulted in a severe heat risk across much of the country and in some instances an extreme heat risk. The majority of locations experienced at least a moderate risk (Figure 2). On 18/07/2022 there were four occasions, each at different stations, where CCI exceeded the threshold for extreme heat risk, on 19/07/2022 there were 22 occasions.

**Figure 2.**
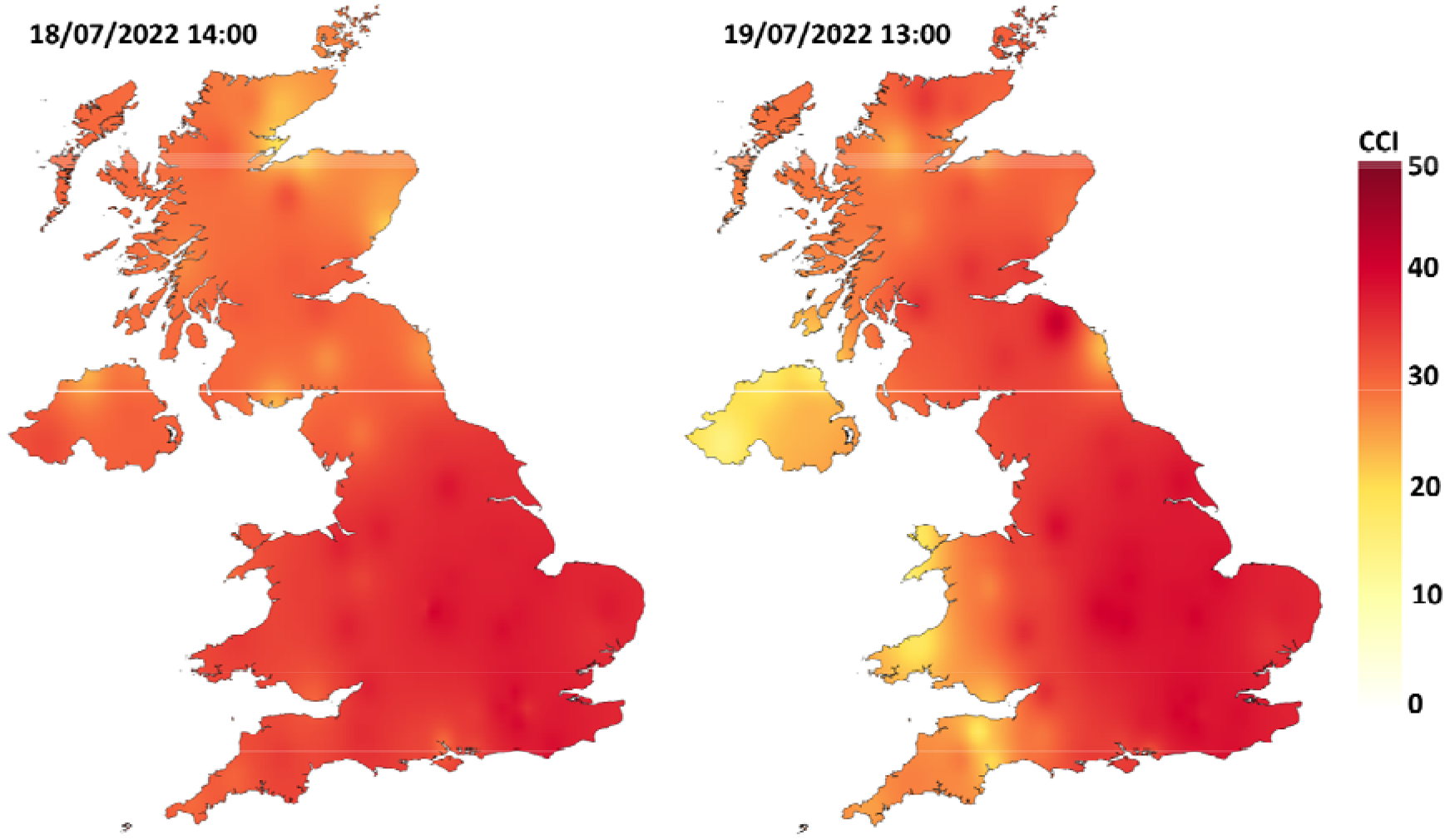
CCI maps for the UK at 14:00 on 18/07/22 and at 13:00 on 19/07/22. Maps are for the period with the highest mean CCI for the given day. Note that data extrapolated to island locations is derived from mainland weather station data.

**Figure 3.**
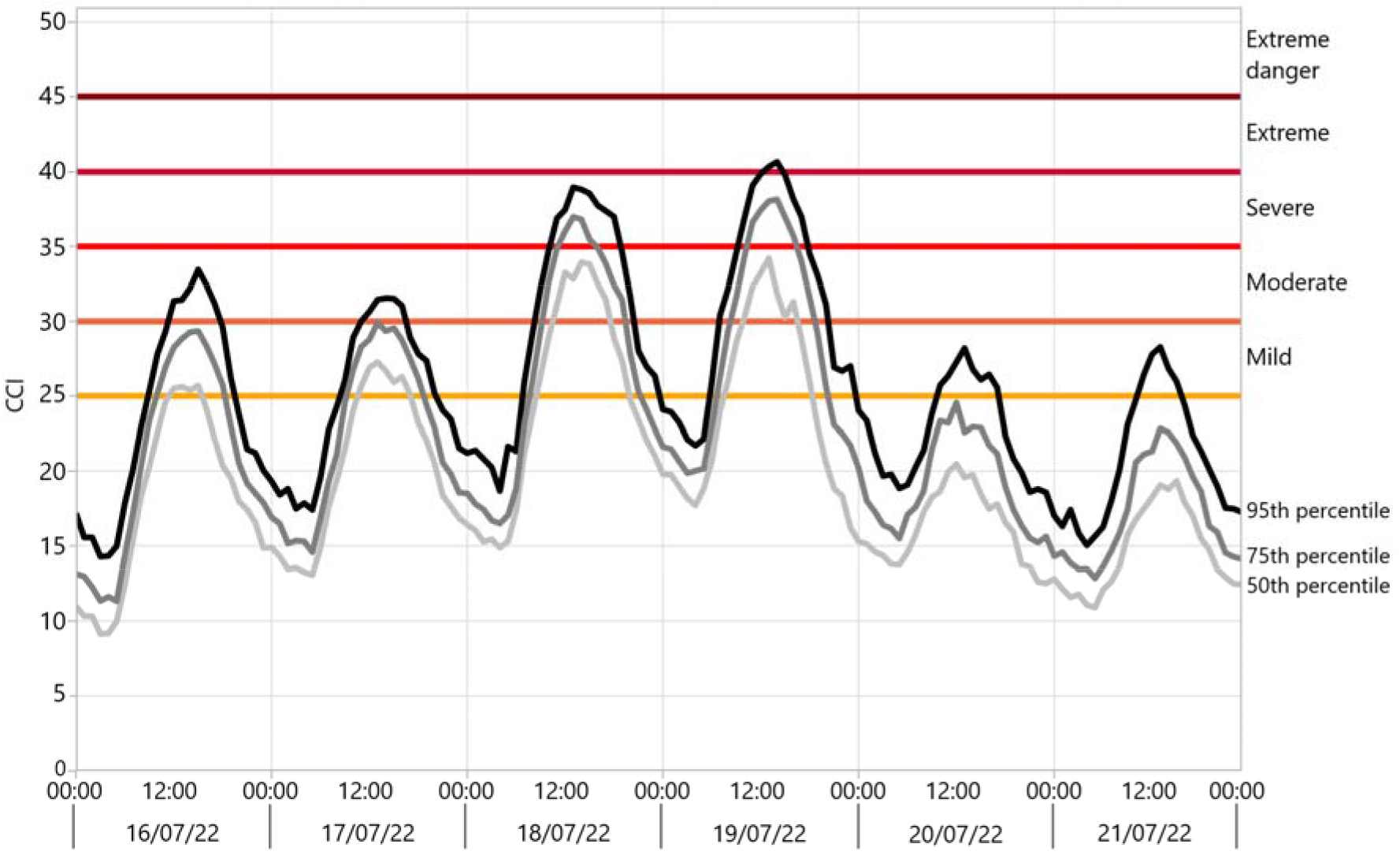
Summary of heat stress risk across the UK from 16/07/22 to 21/07/2022 including 50^th^, 75^th^, and 95^th^ percentiles.

The station with the highest mean CCI value across the two days was in Coventry (SRC_ID 24102); however, this station was in a heavily urbanised area, and thus not typical of livestock systems. Instead, the station with the next highest mean CCI was taken; this was a site (SRC_ID 595) approximately 13 km South-East of Coventry, near Rugby. The site and surrounding area is rural, agricultural in use, with some livestock rearing < 500m from the site – based on satellite imagery taken 16/06/21 (Google, 2021). On the days leading up to 18/07/22, the site experienced weather that posed a moderate heat risk to livestock. On 18/07/22, there was a severe heat stress risk across most of the daytime (Figure 4). This was also the case on 19/07/22, however for a period of approximately 2 hrs CCI thresholds for extreme heat stress risk were exceeded. During the night, between those days, CCI levels remained relatively high only dropping below 25 for a period of a few hours. The two days following the extreme event were far cooler and yielded no apparent heat stress risk.

**Figure 4.**
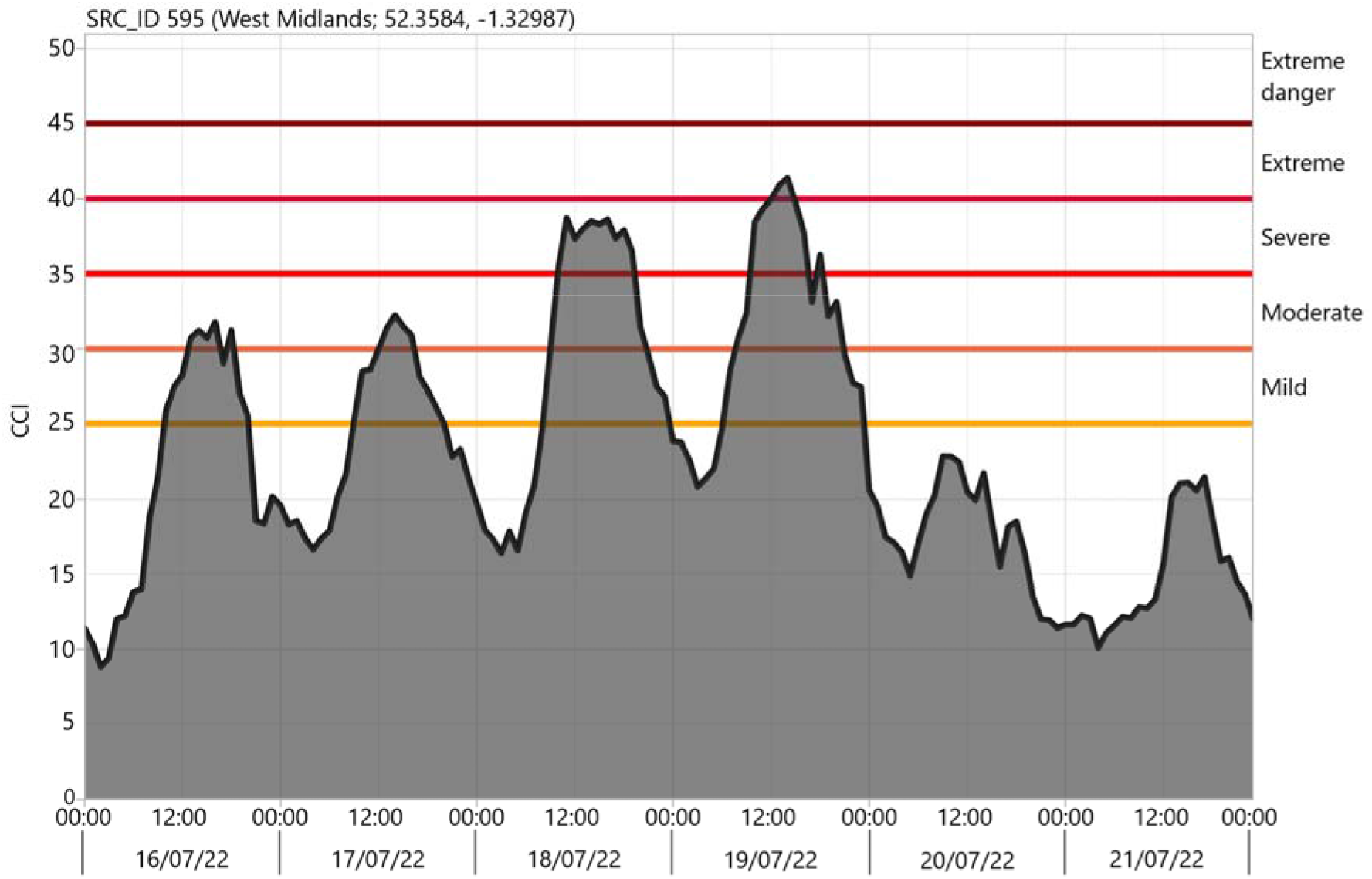
CCI patterns at site SRC_ID 595 during the two days of the “Red Weather Warning” (18/07/22 and 19/07/2022) and two days either side.

The relationship between air temperatue and CCI over the two days was, on average, linear with an approximately 1:1 relationship (Figure 5). However, many individual points yielded air temperature and CCI values that were nearly 10 points out from each other. For example, the point with the highest CCI had a value of 41.2, despite air temperatue being just 33.9°C. The greatest difference between CCI and air tempeature was 8.9 (CCI = 31.4, air temp. = 22.5).

**Figure 5.**
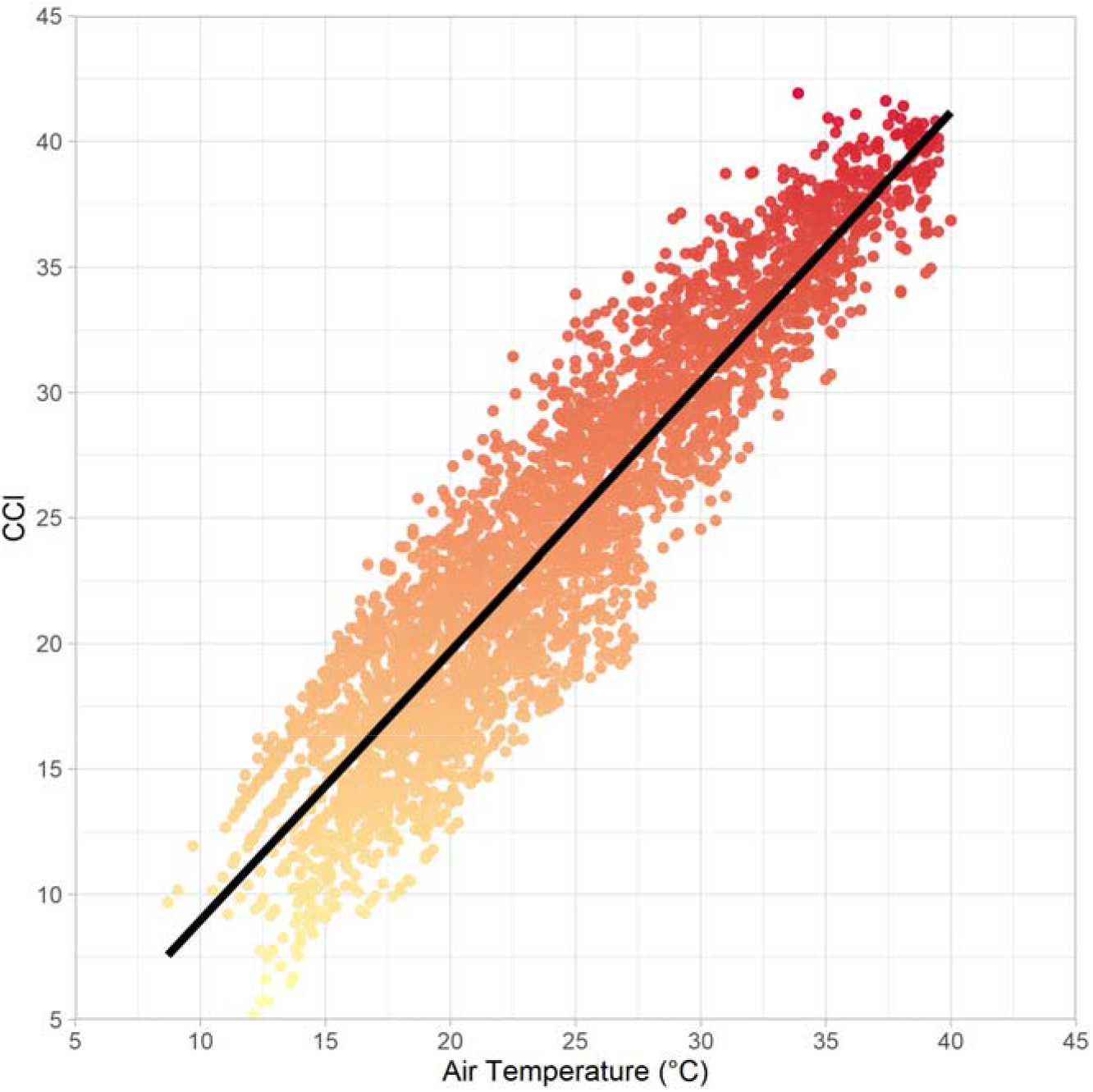
Relationship between CCI and air temperature (°C) across 18/07/22 and 19/07/22. R = 0.86.

Looking at the individual factors that are used to calculate CCI, differences were clear between the extreme heat event of 2022 compared to the two consecutive days in previous Julys with the highest CCI (Figure 6). Air temperature was considerably higher than typical and humidity considerably lower. There appeared to be no large difference in windspeed. The range of solar radiation observed was similar to usual and skewed towards high levels of radiation.

**Figure 6.**
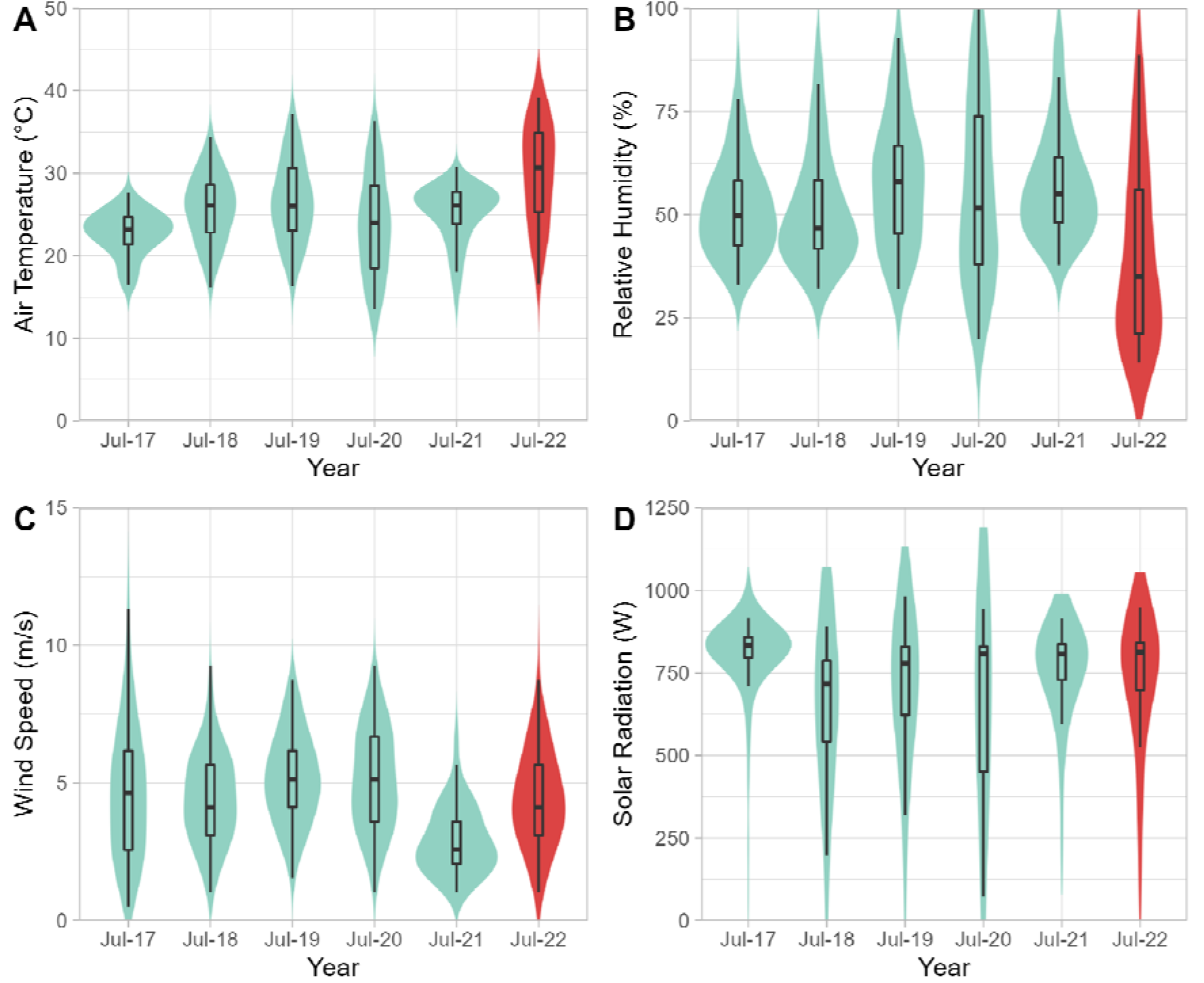
Comparison of weather variables between the extreme heat event of 18/07/22 – 19/07/22 compared to the two consecutive days with the highest CCI of previous Julys (2017-2021). A: Air temperature (°C), B: Relative humidity (%), C: Wind speed (m/s), D: Solar radiation (W). Dates for previous years were: 17-18/07/17, 26-27/07/18, 24-25/07/19, 30-31/07/20, 21-22/07/21.

Satellite images spanning periods before, during and, after the heat event show a clear impact of the weather on vegetation, particularly across the east side of the UK. Normalised Difference Vegetation Index data showed a rapid decline in green vegetation during and after the heat event (Figure 7).

**Figure 7.**
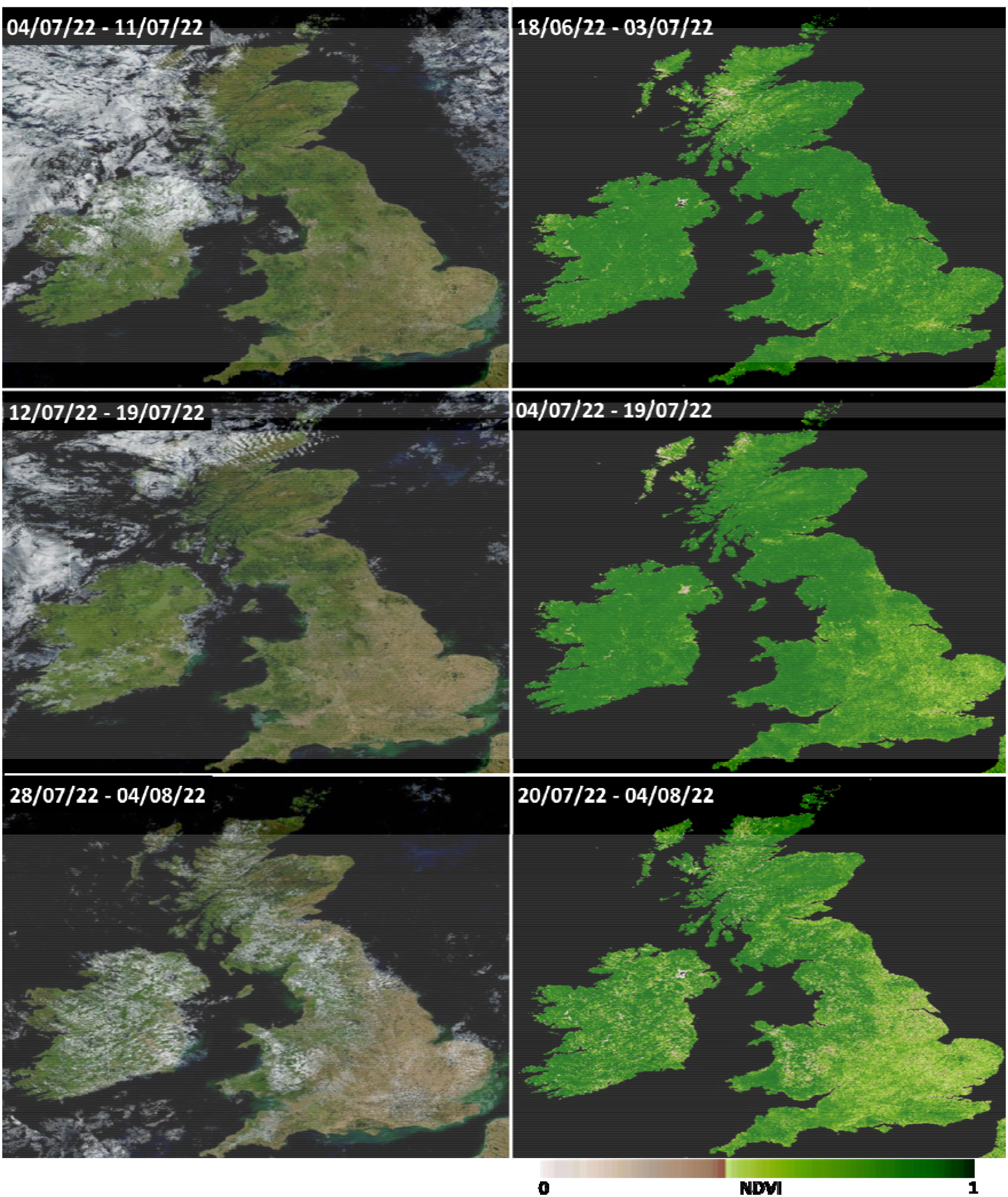
Satellite images taken before, during, and after the period of extreme heat in July 2022. Left side: Land Surface Reflectance (true colour, 8-day composite). Right side: Normalised Difference Vegetation Index (NDVI) (16-day composite). Data originates from Moderate Resolution Imaging Spectroradiometer (MODIS) onboard the Earth Observing System (EOS), obtained via NASA Worldview (NASA, 2022).

The total number of cattle and sheep sent to slaughter (1307 thousand) across the UK in July 2022 was lower (−10.6%) than the mean for the same month on the previous 5 years (1462 thousand) as well as being the lowest across these years (Table 2) (DEFRA, 2022a). This difference appeared to be predominantly due to a reduction in sheep being sent to slaughter (clean sheep -11.7% compared to mean). Milk yield available to dairies for July 2022 was 1176 million litres, representing 16.0% of the year to date. From 2018-2021 the mean of 1182 million litres, representing 16.2% of the year to date. On-fam cattle deaths in July 2022 were also lower than previous years.

**Table 2.** Monthly figures of livestock slaughter, milk yield, and on-farm cattle deaths across the UK for July of years 2017 to 2022 (DEFRA, 2022a). Value in brackets is the proportion (%) that the number represents of the total slaughters/litres in that calendar year from January to July. *milk yield data for July 2017 was removed as reporting methodologies changed between then and July 2018.

|  | 07/2017 | 07/2018 | 07/2019 | 07/2020 | 07/2021 | 07/2022 |
| --- | --- | --- | --- | --- | --- | --- |
| Number of animals slaughtered (thousand head) |  |  |  |  |  |  |
| Steers | 78 (13.3) | 81 (13.6) | 78 (13.4) | 88 (14.9) | 79 (13.8) | 76 (13.6) |
| Heifers | 54 (13.0) | 59 (13.6) | 60 (13.2) | 68 (14.1) | 63 (13.3) | 64 (13.4) |
| Young bulls | 25 (20.2) | 23 (18.9) | 24 (20.3) | 24 (21.1) | 20 (18.0) | 21 (19.1) |
| Cows & Bulls | 52 (14.9) | 60 (15.6) | 53 (13.4) | 60 (15.9) | 51 (14.2) | 52 (14.2) |
| Calves | 7 (10.9) | 7 (11.7) | 9 (12.2) | 5 (9.8) | 5 (14.3) | 5 (11.6) |
| Clean sheep | 1102 (15.7) | 1031 (15.3) | 1090 (15.7) | 1292 (19.4) | 1084 (17.5) | 989 (14.9) |
| Ewes & Rams | 136 (15.3) | 131 (15.1) | 148 (15.7) | 154 (17.6) | 111 (17.5) | 100 (14.7) |
| TOTAL | 1454 (15.4) | 1392 (15.1) | 1462 (15.4) | 1691 (18.5) | 1413 (16.8) | 1307 (14.8) |
| Milk yield (million litres) |  |  |  |  |  |  |
| Cow's milk | * | 1167 (16.3) | 1188 (16.2) | 1187 (16.4) | 1186 (16.0) | 1176 (16.0) |
| On-farm deaths (single head) |  |  |  |  |  |  |
| Cattle | 28,048 | 31,982 | 28,573 | 26,085 | 26,427 | 24,749 |

## 4 Discussion

The heat wave of July 2022 posed a sustained severe, and occasionally extreme, heat stress risk to livestock across many areas of the UK. The effects were felt most in the Midlands and South-East, with other regions suffering to a lesser extent and thus there being a potentially lower risk to animals. Climate modelling predicts that such events are to become more frequent and more extreme due to the effects of climate change (Meehl and Tebaldi, 2004). The impact that heat waves have on livestock depends on variables such as duration (e.g., consecutive days above the critical threshold) and intensity (e.g., level of heat stress reached and number of hours animals are exposed) of the event, and physiological stage, breed, acclimation capacity of the animals. The UK livestock industry needs to invest now and prepare itself for that eventuality to mitigate against the animal welfare, environmental, and economic losses that livestock heat stress can yield.

Although daytime temperatures were high, a reasonable degree of night-time cooling was evident, which is likely to have alleviated overall risk. If this event had lasted into a third or fourth day, that may not have been the case. Night-time cooling may be an effective natural method to alleviate the thermoregulatory limitations of a warm climate (Scott et al., 1983). The ability of cattle to cool (dissipate heat) at night appears to be important for minimizing overall heat load and contributing to the maintenance of normal behaviour and feeding activity (Mader et al., 2006). Cattle that do not cool down at night are prone to achieving greater body temperatures during hot days, whereas cattle that can cool at night can keep peak body temperatures at or near those of cattle that tend to consistently maintain lower body temperatures (Mader et al., 2010a). In moderate-productive dairy cows, cool nights may help to cope with the heat load (Jara et al., 2016); Beede et al. (1993) mention that the night cooling can restore milk production through its effect in restoring dry matter intake. Cool period of less than 21°C for 3 to 6 h will minimize the decline in milk yield (Igono et al., 1992), whereas cows exposed to heat stress for 8 consecutive days show decreased milk fat and protein contents (Ouellet et al., 2019). When using milk yield and mortality risk as indicators, it has been concluded that temperature drops at night below the traditional 72 THI threshold alleviate the effects of heat stress in dairy cows (Nienaber and Hahn, 2007). Regarding duration of heat stress and acclimation capacity of each cow, Galán et al. (2018) performed a systematic review and found that these two factors affect the value of the response; rectal temperature, respiration and heart rates are observed to increase during the early days of exposure but then to drop while the fall in dry matter intake is less severe after three weeks of warm temperatures, suggesting that cows start to acclimate. The duration of the acclimation process (9 to 14 days) varies with breed (Bernabucci et al., 2010). In the case of feed lot cattle, West (2003) found that severe heat waves increase the likelihood for mortality, and several hours of THI > 84 with little or no night-time recovery of THI = 74 can result in the death of vulnerable animals. Thus, global warming could create conditions that not only impair productivity of cattle but increase mortality of cattle in the absence of protective facilities.

Dunn et al. (2014) studied two heat waves that occurred in the UK with the peak temperatures taking place on 10 August 2003 and 19 July 2006 respectively. The authors found that only four herds (out of 17 analysed) showed any indication of a decrease in milk monthly yields during the summers of 2003 and 2006 and suggested that the monthly measurement interval may have masked the impacts as the persistence of any effect of heat stress appears to be low. They reported that there are 0.8 days with THI>70 on average in the UK (over 1973–2012), and during the two years with summer heatwaves this value increased to 2.7 and 2.8 days (2003 and 2006). The authors project that the number of days exceeding the THI threshold for the onset of heat stress (i.e., 70) will increase. For southern parts of the UK this could increase from an average 1–2 per year to over 20 per year by 2100, with correspondingly more during heatwave events.

The reduction in green vegetation appeared to be as a direct consequence of the extreme heat event, hot weather, and low rainfall around that time. The reduction in vegetation availability and production will limit forage dry matter allowance for ruminant herds/flocks. This could lead to associated welfare and economic loses if carrying capacity falls below stocking rate, or if forage quality deteriorates. Grass typically has a high moisture content and in normal conditions a large portion of ruminants water intake is through grass consumption (Minson, 2012). The drying of grass may therefore reduce ruminant water intake, increasing heat stress risk. The ability for livestock to compensate, through voluntary water intake from troughs (or alike) will vary from farm to farm. Furthermore, as ambient temperature increases, so may water intake requirements (Arias and Mader, 2011; Winchester and Morris, 1956). Water intake is typically greatest when water temperatures are warm (Huuskonen et al., 2011; Petersen et al., 2016), however there is a tipping point were water too warm will result in reduced intake (Parish and Karisch, 2022). Having to walk longer distances to obtain water, potentially uphill and out of shade cover, may also contribute towards heat stress risk.

The reason behind the low slaughter numbers and slightly low milk yields compared to the average of previous years is unclear and is not conclusively linked to the heat wave event. Data from individual farms, particularly dairies in the Midlands and South-East, may provide insight into the direct impact of this event at local levels. The deployment of scientific resources in advance of such events in the future would help to better quantify and understand these impacts on UK livestock. This could include digital boluses, thermal imaging, welfare assessments, and physiological and immunological sampling Despite not having such high-resolution data in this instance, it is highly likely that large numbers of livestock suffered welfare losses by means of discomfort, though without long-term impacts. It is also likely that a smaller number of livestock suffer more acute effects resulting in physiological harm.

The location of highest air temperatures did not exactly match up to those with greatest CCI scores – though the two do strongly correlate. This highlights a concern that farmers could inadvertently underestimate the heat stress risk to their cattle, by as much as two or potentially even three risk categories, if they were to rely on air temperature forecasts alone, highlighting the value of considering additional weather variables. Air temperature and CCI differed by as much as 10-15 points. Reporting that focuses on air temperature, typical of mainstream weather reporting, risk farms underestimating the risk to their livestock. There may, therefore, be the need for more tailored reporting for the livestock sector.

The CCI includes air temperature, relative humidity, wind speed and solar radiation, therefore allowing to integrate the multiple environmental factors animals perceive when they graze in the fields. In a systematic review, Galan et al., (2018) found that 86% of the studies use the temperature and humidity together (including THI) as a measure of climate, while 36% of the studies also factor in solar radiation, wind speed or other indices that include them (including CCI). These indices are used especially in studies of pasture systems (66% if studies that include rainfall are also considered). The CCI could be the most promising thermal index to assess heat stress for housed dairy cows (Yan et al., 2021). Dunn et al, (2014) stress that solar radiation implicitly influences the basic THI because THI and solar radiation are positively correlated, whilst wind speeds may be unrepresentative of that experienced by dairy cattle, because wind speeds are more dependent on local topography than are temperature and humidity. On the other hand, Yan et al., (2021) found that the CCI showed a better relationship with the animal-based indicators (i.e., rectal temperature, skin temperature, and eye temperature) of heat stress. CCI has the potential to replace the temperature–humidity index in quantifying the severity of heat stress in dairy cows. It is worth noting that, the thresholds for heat-stress risk are arbitrary (Mader et al., 2010b). The exact risk to livestock is dependent on a variety of factors, such as animal characteristics and acclimation. Notably, the hottest areas during this event were in the Midlands and East, which are by no means typically the warmest places in the UK. The critical thresholds proposed by Mader et al. (2010) for CCI were theoretical and based on beef cattle, that are less sensitive to heat stress than dairy cows. These differences are due to breeds characteristics, production, metabolism, feeding plans, and management systems (Summer et al., 2019). Mader et al. (2010) stressed that CCI has a flexible threshold due to the animals’ susceptibility to environmental factors, previous exposure, age, body condition and isolation. Regardless of the cattle category and the production systems, heat stress impairs primarily animal welfare (Summer et al., 2019).

The risks characterised in this study also highlights the potential risk to livestock that are housed or in transportation. Factors such as orientation, stocking density, materials, and ventilation, can be major contributors to indoor housing and transportation conditions. Whilst there are regulations stating that vehicles must be able to maintain temperatures of 5-30°C, this only applies to journeys in excess of 12 hrs within the UK. In a scenario where temperatures approach closer to 40°C this is likely insufficient, especially if the risk of vehicle or ventilation malfunction is considered. It is advised that future developments be considerate of extreme heat (and cold) in the design of housing facilities and vehicles for any livestock.

### 4.1 Mitigation and intervention

Unlike humans, livestock have no forewarning of weather, no ability to plan for it, and limited capabilities to mitigate it. It is thus duty of their owners and responsible agencies to protect them. With such events predicted to become more probable and more severe, it is important that both short- and long-term strategies are implemented to reduce the heat risk to animals in future events.

The high dry matter intake requirement of ruminants may make the utilisation of shade difficult. Animals may need to break shade cover in order to graze, putting them at increased heat risk. Providing conserved forage (e.g., silage, hay) in shaded areas could reduce the need for animals to leave shade and reduce the energy they have to expend to feed. Converting areas of pasture to silviculture could also address this, by providing an environment that allows cattle to graze with a high level of shade provision, representing a potential synergy between animal welfare and environmental sustainability (Rivero and Lee, 2022). As well as providing shade, tree cover has also been found to reduce ground surface and soil temperatures (Lerman and Contosta, 2019).

Provision of water is essential and water troughs should be placed in accessible areas in or near shade, to prevent cattle overexerting themselves to reach it. Furthermore, water must be prevented from getting too hot as this can exacerbate heat stress. A number of small portable trough solutions (named such as ‘mini’, ‘micro’ or ‘drag’ troughs) are available. These are quick and easy to deploy and in preparation for an extreme heat these can be placed in shaded areas and/or at a high frequency to ensure ease of access and proximity.

Another long-term solution worthy of consideration is the genetic composition of UK livestock and the extent to which animals are suited for a warming climate and extreme heat. This is arguably most important in the context of dairy cattle, due to the high metabolic demand of milk production. There might be a case for including new non-economic traits in the breading objectives for genetic selection of ruminant livestock in the UK, such as “heat tolerance” (Rivero et al., 2021).

The heat experienced in the UK in July 2022 was extreme by UK standards. However, livestock are successfully reared elsewhere in the world in places where such conditions are far more common and often more extreme. Consequently, there may be opportunities for the UK sector to learn from the experience of other countries as the climate warms. Government agencies such as the DEFRA may also wish to consider plans for future heat events that warrant Met Office ‘Red Weather Warnings”, such as restrictions and responsibilities that kick-in over such periods for the protection of livestock (e.g., reducing maximum travel time or pausing travel).

## 5 Conclusion

The record-breaking heat of July 2022 must serve as warning to livestock production in the UK and elsewhere. We cannot know when the next such event will occur, how long it will last, or its intensity. However, we do know that these events will increase in likelihood and severity and whilst we must be wary of knee-jerk reactions, it is also necessary that we prepare today for the world of tomorrow. This will require that systems are designed to minimise heat stress risk were possible, such as through water and shade provision. But it may also mean that mechanisms are in place for such events, such as temporary limitations on transport and movement.

## 6 Acknowledgements

None.

## 9 Ethical Approval

No ethical approval was approved for this study.

## 10 Author Contributions

AC – Concept, study design, data analysis, writing

JR – Study design, data interpretation, writing.

## References

Arias, R.A., Mader, T.L., 2011. Environmental factors affecting daily water intake on cattle finished in feedlots. J. Anim. Sci. 89, 245–251. https://doi.org/10.2527/jas.2010-3014

Beede, D., Bray, R., Bucklin, F., 1993. Planifique su estrategia contra el calor. Venezuela Bov. 7, 38–39.

Bernabucci, U., Lacetera, N., Baumgard, L.H., Rhoads, R.P., Ronchi, B., Nardone, A., 2010. Metabolic and hormonal acclimation to heat stress in domesticated ruminants. Anim. Int. J. Anim. Biosci. 4, 1167–1183. https://doi.org/10.1017/S175173111000090X

Berry, I.L., Shanklin, M.D., and H. D. Johnson, 1964. Dairy Shelter Design Based on Milk Production Decline as Affected by Temperature and Humidity. Trans. ASAE 7, 0329–0331. https://doi.org/10.13031/2013.40772

Carvajal, M.A., Alaniz, A.J., Gutiérrez-Gómez, C., Vergara, P.M., Sejian, V., Bozinovic, F., 2021. Increasing importance of heat stress for cattle farming under future global climate scenarios. Sci. Total Environ. 801, 149661. https://doi.org/10.1016/j.scitotenv.2021.149661

DEFRA, 2022a. Historical statistics notices on the number of cattle, sheep and pigs slaughtered in the UK. Department for Environment, Food and Rural Affairs.

DEFRA, 2022b. Historical national statistics notices on milk utilisation by dairies. Department for Environment, Food and Rural Affairs.

Dunn, R.J.H., Mead, N.E., Willett, K.M., Parker, D.E., 2014. Analysis of heat stress in UK dairy cattle and impact on milk yields. Environ. Res. Lett. 9, 064006. https://doi.org/10.1088/1748-9326/9/6/064006

Galán, E., Llonch, P., Villagrá, A., Levit, H., Pinto, S., Prado, A. del, 2018. A systematic review of non-productivity-related animal-based indicators of heat stress resilience in dairy cattle. PLOS ONE 13, e0206520. https://doi.org/10.1371/journal.pone.0206520

Garner, J.B., Douglas, M., Williams, S.R.O., Wales, W.J., Marett, L.C., DiGiacomo, K., Leury, B.J., Hayes, B.J., Garner, J.B., Douglas, M., Williams, S.R.O., Wales, W.J., Marett, L.C., DiGiacomo, K., Leury, B.J., Hayes, B.J., 2017. Responses of dairy cows to short-term heat stress in controlled-climate chambers. Anim. Prod. Sci. 57, 1233–1241. https://doi.org/10.1071/AN16472

Google, 2021. Google Maps.

Hahn, G., Gaughan, J., Mader, T., Eigenberg, R., 2009. Chapter 5: Thermal Indices and Their Applications for Livestock Environments. Livest. Energ. Therm. Environ. Manag. https://doi.org/10.13031/2013.28298

Huuskonen, A., Tuomisto, L., Kauppinen, R., 2011. Effect of drinking water temperature on water intake and performance of dairy calves. J. Dairy Sci. 94, 2475–2480. https://doi.org/10.3168/jds.2010-3723

Igono, M.O., Bjotvedt, G., Sanford-Crane, H.T., 1992. Environmental profile and critical temperature effects on milk production of Holstein cows in desert climate. Int. J. Biometeorol. 36, 77–87. https://doi.org/10.1007/BF01208917

IPCC, 2022. Climate Change 2022: Impacts, Adaptation, and Vulnerability. Contrib. Work. Group II Sixth Assess. Rep. Intergov. Panel Clim. Change. https://doi.org/10.1017/9781009325844.

Jara, I.E., Keim, J.P., Arias, R.A., 2016. Behaviour, tympanic temperature and performance of dairy cows during summer season in southern Chile. Arch. Med. Vet. 48, 113–118. https://doi.org/10.4067/S0301-732x2016000100014

Lees, A.M., Sejian, V., Wallage, A.L., Steel, C.C., Mader, T.L., Lees, J.C., Gaughan, J.B., 2019. The Impact of Heat Load on Cattle. Animals 9, 322. https://doi.org/10.3390/ani9060322

Lerman, S., Contosta, A., 2019. Lawn mowing frequency and its effects on biogenic and anthropogenic carbon dioxide emissions. Landsc. Urban Plan. 182, 114–123. https://doi.org/10.1016/j.landurbplan.2018.10.016

Mader, T.L., Davis, M.S., Brown-Brandl, T., 2006. Environmental factors influencing heat stress in feedlot cattle. J. Anim. Sci. 84, 712–719. https://doi.org/10.2527/2006.843712x

Mader, T.L., Gaughan, J.B., Johnson, L.J., Hahn, G.L., 2010a. Tympanic temperature in confined beef cattle exposed to excessive heat load. Int. J. Biometeorol. 54, 629–635. https://doi.org/10.1007/s00484-009-0229-0

Mader, T.L., Johnson, L.J., Gaughan, J.B., 2010b. A comprehensive index for assessing environmental stress in animals. J. Anim. Sci. 88, 2153–2165. https://doi.org/10.2527/jas.2009-2586

Meehl, G.A., Tebaldi, C., 2004. More Intense, More Frequent, and Longer Lasting Heat Waves in the 21st Century. Science 305, 994–997. https://doi.org/10.1126/science.1098704

Met Office, 2022. UK prepares for historic hot spell [WWW Document]. Met Off. URL https://www.metoffice.gov.uk/about-us/press-office/news/weather-and-climate/2022/red-extreme-heat-warning (accessed 7.27.22).

Met Office, 2012. Met Office (2012): Met Office Integrated Data Archive System (MIDAS) Land and Marine Surface Stations Data (1853-current) [WWW Document]. NCAS Br. Atmospheric Data Cent. URL http://catalogue.ceda.ac.uk/uuid/220a65615218d5c9cc9e4785a3234bd0

Minson, D., 2012. Forage in Ruminant Nutrition. Elsevier.

NASA, 2022. NASA Worldview [WWW Document]. URL https://worldview.earthdata.nasa.gov/

Ouellet, V., Cabrera, V.E., Fadul-Pacheco, L., Charbonneau, É., 2019. The relationship between the number of consecutive days with heat stress and milk production of Holstein dairy cows raised in a humid continental climate. J. Dairy Sci. 102, 8537–8545. https://doi.org/10.3168/jds.2018-16060

Parish, J., Karisch, B., 2022. Beef Cattle Water Requirements and Source Management (No. P2490). Mississippi State University.

Petersen, M.K., Muscha, J.M., Mulliniks, J.T., Roberts, A.J., 2016. Water temperature impacts water consumption by range cattle in winter1. J. Anim. Sci. 94, 4297–4306. https://doi.org/10.2527/jas.2015-0155

QGIS, 2022. QGIS Geographic Information System.R Core Team, 2021. R: A language and environment for statistical computing. R Studio Team, 2020. RStudio: Integrated Development for R. RStudio.

Rivero, M.J., Eisler, M. C., Takahashi, T., Lee, M.R.F., 2021. Key traits for ruminant livestock across diverse production systems in the context of climate change: perspectives from a global platform of research farms. Reprod. Fertil. Dev. 33, 1–19. https://doi.org/10.1071/RD20205

Rivero, M.J., Lee, M.R.F., 2022. A perspective on animal welfare of grazing ruminants and its relationship with sustainability. Anim. Prod. Sci. https://doi.org/10.1071/AN21516

Scott, I.M., Johnson, H.D., Hahn, G.L., 1983. Effect of programmed diurnal temperature cycles on plasma thyroxine level, body temperature, and feed intake of holstein dairy cows. Int. J. Biometeorol. 27, 47–62. https://doi.org/10.1007/BF02186300

Summer, A., Lora, I., Formaggioni, P., Gottardo, F., 2019. Impact of heat stress on milk and meat production. Anim. Front. 9, 39–46. https://doi.org/10.1093/af/vfy026

Urbanek, S., Horner, J., 2020. Cairo: R Graphics Device using Cairo Graphics Library for Creating High-Quality Bitmap (PNG, JPEG, TIFF), Vector (PDF, SVG, PostScript) and Display (X11 and Win32) Output.

West, J.W., 2003. Effects of heat-stress on production in dairy cattle. J. Dairy Sci. 86, 2131–2144. https://doi.org/10.3168/jds.S0022-0302(03)73803-X

Wickham, H., 2016. ggplot2: Elegant Graphics for Data Analysis.

Winchester, C.F., Morris, M.J., 1956. Water Intake Rates of Cattle. J. Anim. Sci. 15, 722–740. https://doi.org/10.2527/jas1956.153722x

Yan, G., Li, H., Shi, Z., 2021. Evaluation of Thermal Indices as the Indicators of Heat Stress in Dairy Cows in a Temperate Climate. Anim. Open Access J. MDPI 11, 2459. https://doi.org/10.3390/ani11082459

